# A General Framework for Branch Length Estimation in Ancestral Recombination Graphs

**DOI:** 10.1101/2025.02.14.638385

**Authors:** Yun Deng, William S. DeWitt, Yun S. Song, Rasmus Nielsen

**Affiliations:** Center for Computational Biology, University of California, Berkeley, USA; Computer Science Division, University of California, Berkeley, USA; Department of Genome Sciences, University of Washington, Seattle, WA, USA; Department of Statistics, University of California, Berkeley, USA; Department of Integrative Biology, University of California, Berkeley, USA; Center for GeoGenetics, University of Copenhagen, Denmark

**Keywords:** Ancestral Recombination Graph, branch length estimation, uninformative prior

## Abstract

Inference of Ancestral Recombination Graphs (ARGs) is of central interest in the analysis of genomic variation. ARGs can be specified in terms of topologies and coalescence times. The coalescence times are usually estimated using an informative prior derived from coalescent theory, but this may generate biased estimates and can also complicate downstream inferences based on ARGs. Here we introduce, POLEGON, a novel approach for estimating branch lengths for ARGs which uses an uninformative prior. Using extensive simulations, we show that this method provides improved estimates of coalescence times and lead to more accurate inferences of effective population sizes under a wide range of demographic assumptions (population expansion, bottleneck, split, etc). It also improves other downstream inferences including estimates of mutation rates. We apply the method to data from the 1000 Genomes Project to investigate population size histories and differential mutation signatures across populations. We also estimate coalescence times in the HLA region, and show that they exceed 30 million years in multiple segments.

**Significance Statement:** Model misspecification is a common challenge in population genetic inference, as oversimplified mathematical models often fail to capture complex evolutionary processes. Here we introduce a novel framework for branch length estimation in whole-genome genealogies, which enables accurate inference using non-informative priors and subsequent posterior calibration, rather than relying on heavily parametrized models. This flexible approach has been validated in various simulation settings. When applied to real genomic data, it enables robust inference of demography, mutation rates, and transspecies polymorphisms.

---

**T**he genetic relationships between the DNA sequences in a sample can be described by a tree. However, in the presence of recombination, the trees might differ from position to position in the genome (1). The set of all trees in the genome, and corresponding change points (Figure 1A), is represented by the Ancestral Recombination Graph (ARG) (1–5). The generative ancestral process creating ARGs is the coalescent with recombination (2). Simulations of ARGs under this model is relatively straightforward (6, 7), but inferring ARGs from DNA sequence data is known as a notoriously difficult problem (8, 9). The difficulty of ARG inference is mainly caused by the high dimensionality of the ARG; there are a lot of possible topologies for each local tree and the genome might contain a large number of such trees. Every recombination event will generate a new tree that might have a new topology. Even when topologies do not change after recombination, branch lengths might change (10) and ARG inference contains a dual problem of inferring both topology and branch lengths.

**Fig. 1.**
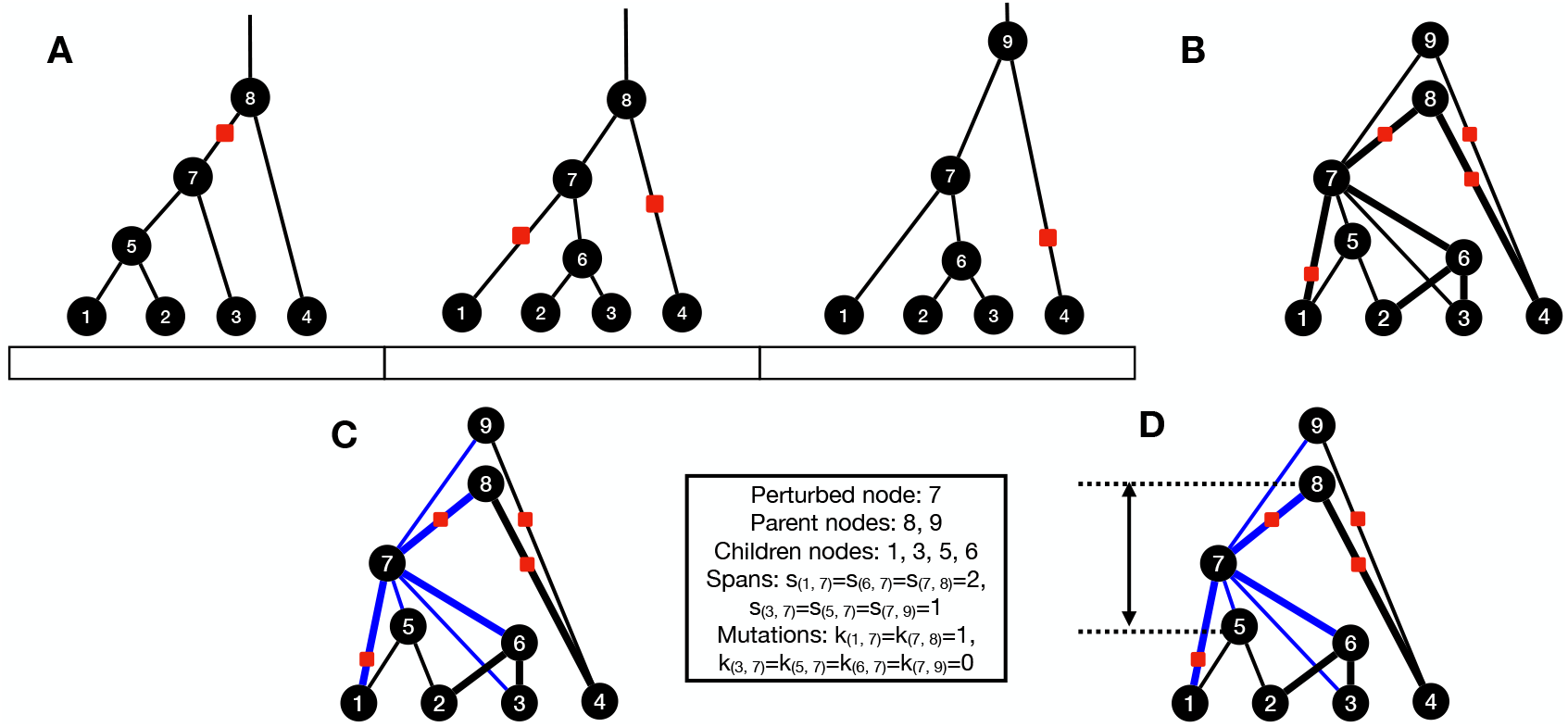
Methodology overview of POLEGON. The tree sequence format of local genealogies on each non-recombining block (A) can be converted to a DAG (B), by merging the nodes with the same index across trees. The width of the edges indicates their spatial spans in the tree sequence. When updating the age of a particular node (C), its age can only move within the interval (D) determined by Eq. (2), as the blue branches must have positive branch length. The age of the node is perturbed in this interval with an MCMC algorithm to sample from Eq. (3). We denote the number of mutations mapped to the branch connecting node *u* and *v* by *k*_(*u,v*)_, and the total length of the genomic segments where node *u* and *v* are adjacent by *s*_(*u,v*)_, where *u* and *v* are unordered.

One important feature of ARGs is the “node persistence” property, i.e. that a node can be shared by multiple marginal trees. For example, in Figure 1A, all three trees share node 7. Each node in the ARG represents a genomic segment of ancestral genetic material, which might span over multiple local trees. Node persistence reduces the dimensionality of the ARG inference problem and explains how a tree in one position of the genome is informed by trees in adjacent segments of the genome. This property also facilitates the efficient storage and simulation of ARGs implemented in the package tskit, which leverages this property in its “tree sequence” format (7, 11).

There are many methods for estimating ARGs including ARGweaver (9), Relate (12), tsinfer (13), KwARG (14), ARG-Needle (15), and SINGER (16). Several of these methods require a branch length estimation step after the topology has been inferred. Relate (12) can estimate branch lengths for each locally inferred topology in the genome under a specified demographic model, tsdate (17) dates the node ages in the inferred ARG topology from tsinfer (13), and ARG-Needle normalizes node ages to ensure that they match a distribution obtained using simulations under a given demography model. To our knowledge, only Relate can estimate branch length and the population size history jointly. The “EstimatePopulationSize.sh” module in Relate achieves this using a Markov Chain Monte Carlo (MCMC) algorithm to re-estimate branch length given a demographic prior and subsequently estimates a new demographic model from the new branch length. Relate alternates between updating branch length and the demography model similarly to an EM-algorithm until convergence is achieved (the default is 10 iterations). This alternation between branch length and demography estimation is computationally demanding and will, in most applications, be the computational bottleneck in the inferences.

The use of a coalescent prior in previous work is a natural approach for estimating coalescence time (or equivalently branch lengths) in ARGs. However, it may bias the estimates when the assumed coalescent prior is mis-specified, which typically will be the case. The true distribution of coalescence times in natural populations is highly complicated and depends on the history of population sizes and population structure. Here we introduce a novel approach, POLEGON (**P**rior-**O**blivious **L**ength **E**stimation in **G**enealogies with **O**riented **N**etworks) for estimating branch lengths for ARGs using an uninformative uniform prior for coalescence times, which currently works for ARG topology inferred by SINGER (16). POLEGON can also be applied to other ARG topology inference methods, though specific refinements of inferred ARGs might be needed to further improve results (for example, treatment of polytomies in ARG topologies inferred by tsinfer). Using simulations, we show that it provides improved estimates of coalescence times under a wide range of demographic assumptions.

## Results

### Methodology overview

An ARG can be converted to a directed acyclic graph (DAG) of only the coalescence nodes (without recombination nodes), by identifying the nodes with the same index across local trees, and the branches with the same child and parent node (Figure 1B). This representation was introduced as the “genome-ARG” in (4). We note that branch length estimation is equivalent to node age estimation here, because the length of a branch is the age difference of the parent and child nodes associated with the branch. In the following, we will describe our new method in terms of node age estimation in the DAG.

To estimate the node ages, we will use an MCMC algorithm algorithm that iteratively updates the marginal node ages node by node, while the ages of all other nodes are fixed. Let nodes *i* and *n* be adjacent nodes (i.e., there is an edge between them) in the DAG, then we define the span, *s*_(*i,n*)_, as the total length of the genomic segments where *i* and *n* are adjacent (Figure 1C). We assume that mutations can be mapped on edges in the ARG under the assumption of the infinite-sites model, using a parsimony criterion (that minimizes the number of mutations needed to explain the observed site patterns), or using some other algorithm, such as maximum likelihood under a given mutation rate matrix (see e.g. (12, 13, 16)). We denote the number of mutations mapped to the branch connecting node *i* and *n* by *k*_(*i,n*)_, where *i* and *n* are unordered. The age of node *n* is denoted by *t*_*n*_. For each edge, *i* adjacent to node *n*, we then assume that *k*_(*i,n*)_ follows a Poisson distribution:

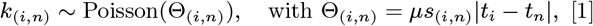

where *µ* is the mutation rate, i.e., the expected number of mutations per site per time unit. Note that *t*_*n*_ is not a completely free parameter; it is constrained by the ages of its children nodes and its parent nodes (Figure 1D). Let *l*_*n*_ be the age of node *n*’s oldest child node and *u*_*n*_ be the age of node *n*’s youngest parent node. Then,

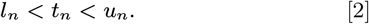

Therefore, we define the marginal likelihood of the age of node *n*, conditionally on it being in the interval (*l*_*n*_, *u*_*n*_), as:

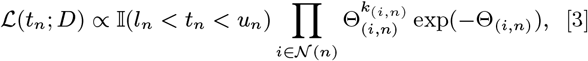

where 𝕀(*l*_*n*_ *< t*_*n*_ *< u*_*n*_) is the indicator function of the condition in (2), (*n*) is the set of all neighbors of node *n*, and *D* is the vector of the number of mutation counts on each branch of the ARG. We use an improper uniform prior (from 0 to ∞) on the node ages, so the marginal posterior is proportional to (3). We note this is equivalent to assuming the coalescent times follow an improper uniform distribution (regardless of the number of lineages), in a single coalescent tree and the full ARG (see *SI Appendix, SI Text*, Section B for detailed discussion). So this avoids the usage of informative coalescent priors as in previous efforts such as Relate and tsdate (12, 17). The informative coalescent priors are right-skewed and their density functions decrease with time (such as exponential distribution for pairwise coalescence time), while the uninformative prior is flat for all times.

We use a Metropolis-Hastings algorithm to iteratively sample from this posterior for all nodes in the graph marginally (Section). When processing the MCMC output, it will often be convenient to take an average over MCMC samples, and it is convenient in this context that the posterior sample averages of the ages of the nodes will satisfy the DAG constraints from Eq. (2). This is because each set of node ages in one posterior sample satisfies the constraints and their averages, as linear combinations, will consequently satisfy the constraints as well.

If the true coalescence process follows the coalescent with recombination (2), or any other common coalescent process, the improper uniform prior on node ages is a misspecified prior, in the sense that it does not match the known/assumed generative process. How to address the problem of prior misspecification is an active topic in Bayesian statistics, and there has been substantial recent research in this area, including solutions based on empirical Bayes methods (18), the Saerens-Latinne-Decaestecker algorithm (19), EM algorithms (20) and more (21, 22). In the current context, the ARG rescaling technique introduced in SINGER (16) can help correct the posterior for misspecification of the prior distribution. Briefly, this rescaling technique relies on the assumption of a constant mutation rate through time, which can be used to define a monotonic transformation of the node ages such that the mutation rate is estimated to be constant from the ARG with adjusted node ages. As such, ARG rescaling attempts to bring coalescence times closer to the posterior distribution that would have been observed, if the true coalescent process had been used as a prior for the node ages (Figure 2A). A caveat of this approach is that it assumes that the overall mutation rate remains constant over time, and we note that mutation rates have been argued to change substantially over time (23). The full algorithm of POLEGON includes such an ARG rescaling step applied to the samples obtained from the MCMC chain (*SI Appendix, SI Text*, Section A).

**Fig. 2.**
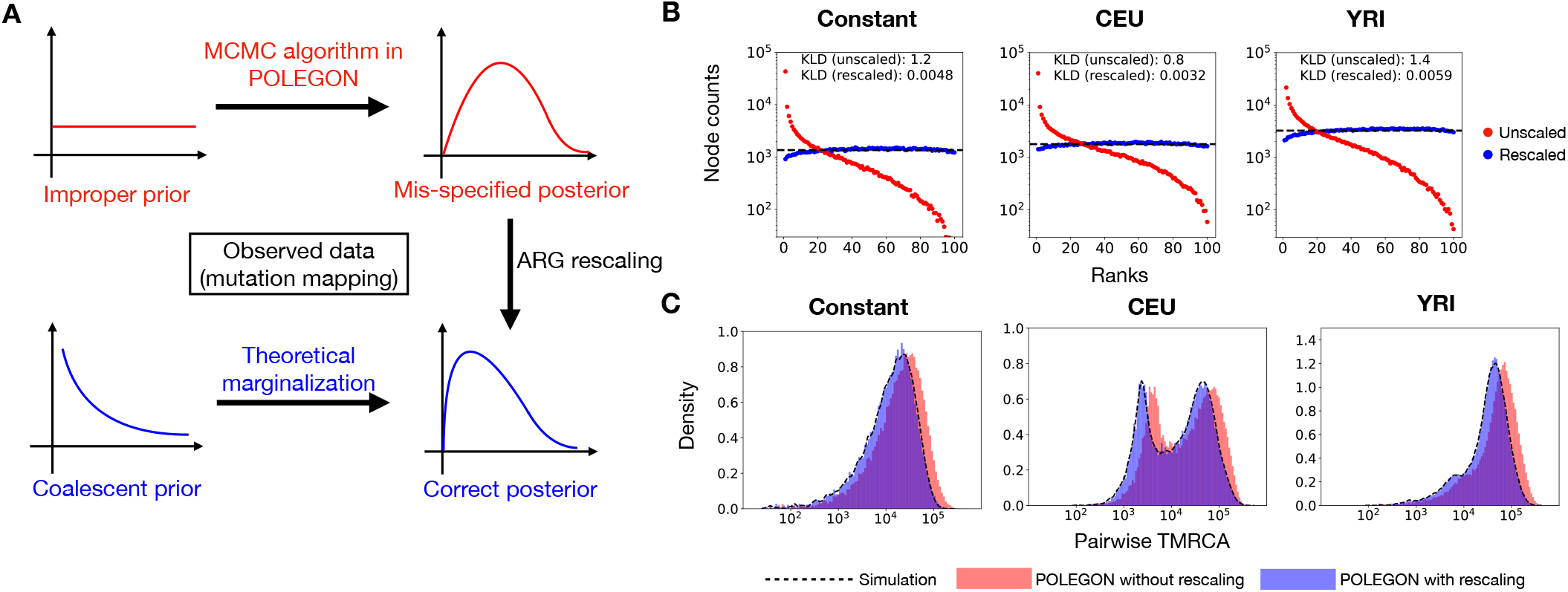
Calibration of distribution with ARG rescaling. (A) ARG rescaling transforms the original mis-specified posterior distribution closer to the correct distribution, even if the true coalescent prior is unknown; (B) The rank plot of the node ages against the node age samples from POLEGON, before (red) and after (blue) rescaling, under simulation under three different demography models: constant size, CEU model and YRI model; (C) The pairwise TMRCA distribution in simulation (black) compared to inferred with POLEGON, with (blue) and without (red) the ARG rescaling operation.

It is worth noting that ARG rescaling does not change the relative order of nodes in the DAG, even when multiple orderings are permitted by the topology. For example, node 5 in Figure 1B can be older or younger than node 6 without violating the DAG constraints. The MCMC algorithm (Section) samples possible relative orderings, while the rescaling step assigns ages given the ordering of nodes.

Relate (12) also provides an MCMC method for obtaining a posterior for the node ages of an ARG. However, Relate only enforces constraints for a marginal coalescent tree given the nodes in that tree, i.e. it does not use the node-persistence property of the ARG. In contrast, POLEGON enforces constraints from the full DAG. Another major difference is in the choice of prior, with Relate enforcing a demographic prior, while POLEGON is using an improper uniform prior with subsequent posterior rescaling. This increases the speed of the program as joint estimation of both demographic parameters and node ages is not necessary, and makes POLEGON more robust and flexible in terms of demographic models it can accommodate. Finally, POLEGON can accommodate spatial variation in mutation rate along the length of the genome.

In terms of demography inference, Relate infers ARG-based population size history by alternatively estimating branch length or coalescence rates while fixing the other (by default using 10 iterations). Here, we estimate the population size history based on pairwise TMRCA distribution, directly after estimating branch lengths with POLEGON, without having to re-estimate branch lengths given the demography model (details in Section). The full details of POLEGON’s algorithm can be found at *SI Appendix, SI Text*, Section A.

### Simulation benchmarks

To benchmark the performance of POLEGON, we carry out simulations using msprime (7), with parameters that are realistic for humans, *µ* = 1.2 *×* 10^−8^, *r* = 1.2 *×* 10^−8^, *L* = 10 Mb and with a sample size of 100 genomes, for 10 replicates. We simulate under three models: (1) constant-size (*N*_*e*_ = 10^4^) and (2) CEU and (3) YRI models from SMC++ (details in *SI Appendix, SI Text*, Section A).

To demonstrate how the combination of uninformative prior and ARG rescaling (Figure 2A) works, we first analytically solve the pairwise coalescent case with unknown constant population size (*SI Appendix, SI Text*, Section C). In this case, we show analytically that the rescaled posterior distribution, in expectation, is exactly identical to the correct distribution (*SI Appendix, SI Text*, Section C), even when the population size history is unknown.

For more complex generative processes of the coalescent with recombination, we use simulations to demonstrate that the ARG rescaling provides well-calibrated posterior distributions. We simulate under the aforementioned three different models, constant *N*_*e*_, a CEU model, and a YRI model (details in *SI Appendix, SI Text*, Section A). We provide POLEGON with the true topology from the simulations, and use a rank plot to diagnose the quality of the posterior samples (24, 25). If the posterior sampling is perfect, then the relative rank of the true node age among the age samples, when simulating data from the prior, should be uniformly distributed between 0 and *M*, where *M* is the number of samples from the MCMC chain (24). This is, to a close approximation, indeed the case for all three different generative models, as evidenced by a low Kullback-Leibler divergence between the empirical rank distribution and a uniform distribution (Figure 2B). Similarly, the estimates of the distribution of pairwise coalescence times from POLEGON match the true distributions well, but only after ARG rescaling (Figure 2C, *SI Appendix, SI Text*, Table S2). We also examined the same demographic models but with *µ/ρ* = 0.25 or 4, to show that this conclusion generalizes to other parameter choices (*SI Appendix, SI Text*, Figure S2).

We compare the inference accuracy of SINGER+POLEGON to that of SINGER and Relate, to evaluate the accuracy gain from additionally using POLEGON. SINGER has previously been shown to provide more accurate estimates in some aspects than other ARG inference methods (16). In the comparisons, we examine the inference accuracy of pairwise time to the most recent common ancestor (pairwise TMRCA), the distribution of pairwise TMRCA, local diversity (the average pairwise TMRCA in genome windows), and local mutation density (the average tree branch length in genome windows). These are statistics similar to those used to evaluate ARG inference performance in (16). In all simulations, we first run SINGER to infer the ARG topology and we then run POLEGON on the MCMC samples from SINGER. Posterior averages of these statistics among samples were calculated and compared to ground truth to evaluate accuracy. The details of the simulation benchmarks can be found at *SI Appendix, SI Text*, Section E.

We compute the local diversity and local mutation density in 10 kb windows, following the branch-length-based definition of (26) (using “diversity” and “segregating sites” API in tskit). We used the mean squared error (MSE) to characterize the inference accuracy for all statistics. For evaluating the distribution of pairwise TMRCAs, we categorize the coalescence times into 20 bins so that each bin occupies 5% of the probability mass in the ground truth distribution, and then compute the symmetrized Kullback-Leibler divergence (KLD), also known as the Jeffreys divergence, between the simulated and inferred distribution with the discretized distributions. For all four tasks, POLEGON+SINGER improves the accuracy over that of SINGER alone, which is itself better than Relate (Figure 3). Importantly, the pairwise TMRCA distribution is more accurately inferred by SINGER+POLEGON than using SINGER alone (Figure 3B), which will lead to improved inferences of demographic history.

**Fig. 3.**
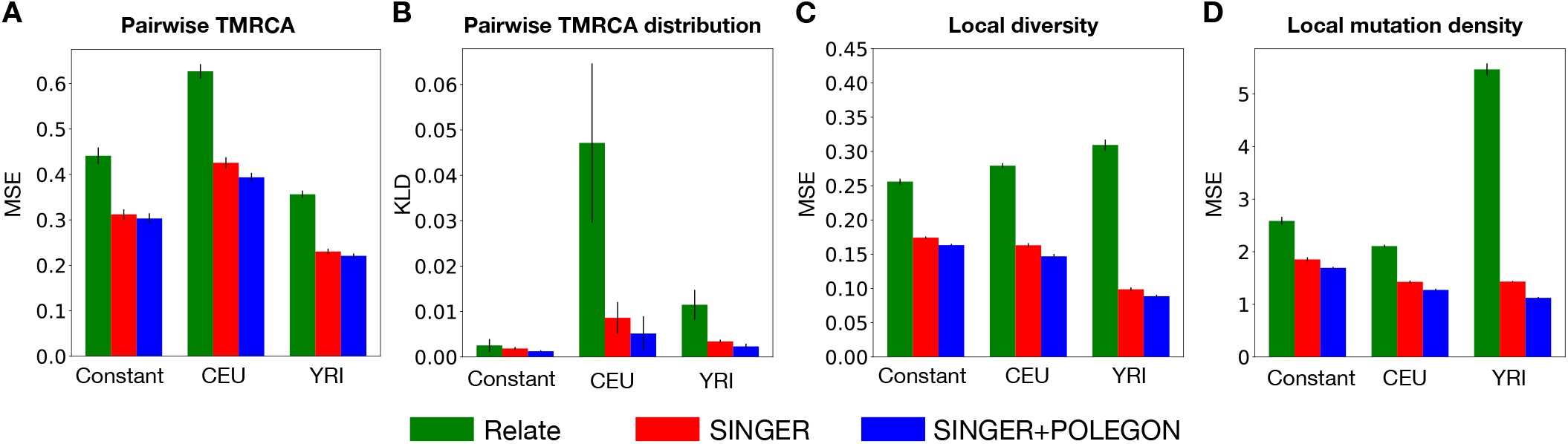
The inference accuracy comparison between Relate (green), SINGER (red) and SINGER+POLEGON (blue), in different aspects: pairwise TMRCA (A), pairwise TMRCA distribution (B), local diversity (C), and local mutation density (D). Error bars indicate standard deviations from 10 replicates.

In real data analyses, the underlying demographic model is likely different from a constant-size panmictic population model, and the demographic model typically needs to be inferred from the data. There exist many demographic inference methods based on different frameworks, for example using the Site Frequency Spectrum (SFS) (27–29), the Sequentially Markovian Coalescent (SMC) for pairs of genomes (30–32), etc. Each of these methods have drawbacks. Reducing population genomic data to an SFS leads to a substantial loss of information, as all linkage/haplotype information is lost. Classical SMC methods typically only handle 2 to 8 genomes (30–32). ARGs provides a powerful alternative for estimating population size changes (12), if the node ages can be estimated accurately. Here we compare the demography inference results from SINGER+POLEGON with other competing methods including MSMC2 (31, 32) (an SMC based method), Relate (12) (based on ARGs), and FitCoal (33) (an SFS based method). We find that, the accuracy of POLEGON is at least comparable to MSMC2 and Relate, and better than that of FitCoal, especially on the resolution of the change history (Figure 4A, B, *SI Appendix, SI Text*, Figure S7). A likely reason is that FitCoal only accommodates a few changes in *N*_*e*_ (it assumes a piece-wise exponential growth/decline), while other methods typically allow for more change points. We also note that using the full information in the ARG as opposed to only a few pairs of sequences can lead to more stable estimates, as indicated by *SI Appendix, SI Text*, Figure S3.

**Fig. 4.**
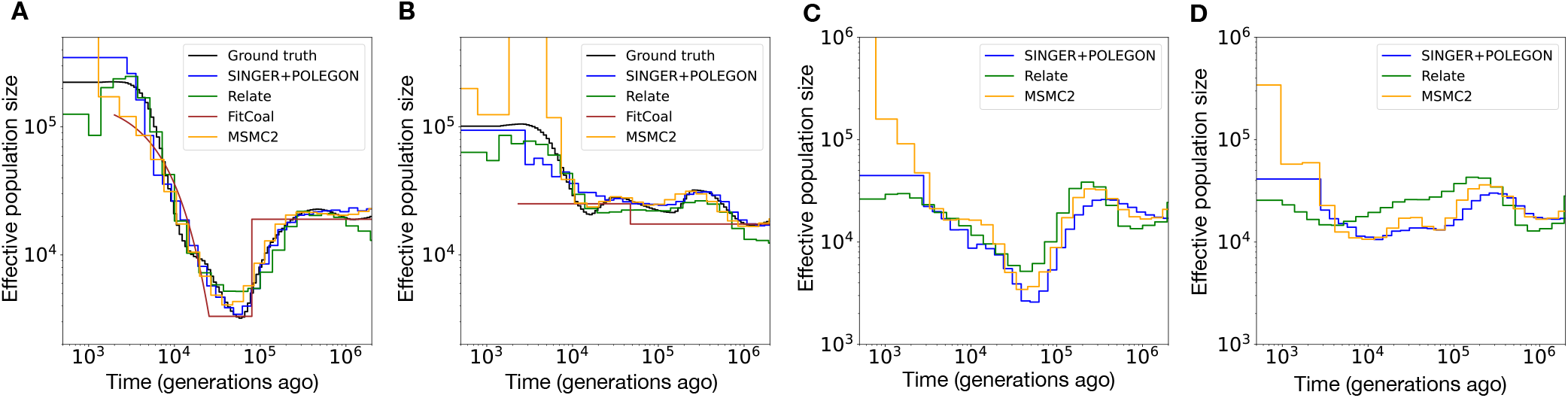
Demography inference results. Simulation benchmark of demography inference with the CEU model (A) and the synthetic model (B), with the inference error from each method provided in the brackets with coalescence rate divergence. We also inferred the population size history for CEU (C) and YRI (D) in the 1000 Genomes Project.

In order to evaluate the performance under a structured demography model, we additionally simulate a model with two populations that split *t* = 10, 000 generations ago (*SI Appendix, SI Text*, Section C), and compare the performance of SINGER+POLEGON to that of Relate in terms of accuracy of inferred cross-population coalescence rates. We find that SINGER+POLEGON infers the cross-coalescence rate dropping towards zero slightly faster after the true divergence time than Relate (*SI Appendix, SI Text*, Figure S4).

### Population size history in YRI and CEU

Non-African populations share an bottleneck roughly 60kya (31) due to the “Out-Of-Africa” migration, which is not shared by African populations. However, it has recently been proposed that African populations experienced a rather extreme bottleneck around 900kya (33). This signal has only been identified by FitCoal but not other methods, such as SMC++, PSMC, MSMC, Relate and so on. Efforts to reproduce their results with other methods also failed (34, 35). More recently, it was argued that the severe bottleneck is a statistical artifact (36). We applied SINGER to the CEU and YRI samples in the 1000 Genomes Project high coverage dataset (37) to obtain estimates of ARG topology. We then used POLEGON to estimate branch lengths and population size history (details in *SI Appendix, SI Text*, Section A). We compared our results to those inferred with MSMC2 (31, 32) and Relate (12), and none of them shows evidence of the sudden, severe bottleneck.

All methods are able to infer the Out-of-Africa bottleneck in CEU and its absence in YRI. They also all observed a mild ancient bottleneck around 1 million years ago (Figure 4C, D). Importantly, this mild bottleneck is also detected in both CEU and YRI, which is expected that it predates the divergence time of CEU and YRI. (36) also reported that the severe bottleneck is likely a statistical artifact and that FitCoal tends to infer a sharp bottleneck in the presence of only mild reductions of population size. We also note that the estimates of population size at very recent times tend to be noisy and unstable, as pairwise coalescences are sparse at this resolution.

### Differential mutation signatures in YRI and CEU

Several different approaches have shown that human populations differ in certain mutation-type specific mutation rates (12, 38– 41). Most notably, a signature of increased mutation rate of TCC to TTC in European populations has been identified using many different approaches (12, 38, 40, 41).

Some methods for estimating mutation rate trajectories are based on allele age estimation from inferred genealogies, and as such, the quality of inferred genealogies and their branch lengths has substantial impact on allele age estimates (12). Additionally, as mutations can only be dated to a specific branch in the ARG, the associated allele age is often assumed to be uniformly distributed along the length of that branch. However, this is only valid under a constant mutation rate model, and it does not apply to the case of temporally varying mutation rates. Therefore, rate estimation based on a uniform distribution on branches tends to be inaccurate if there truly is rate variation over time.

To illustrate this point, we performed a simulation with an elevated mutation rate from 300 to 3,000 generations ago (details in *SI Appendix, SI Text*, Section D). We note here that we fix the node ages from the simulation and only infer the mutation rate trajectory. Even when using the ground truth topology and mutation mappings from simulations, the inferred mutation rate trajectory underestimates the true temporal rate heterogeneity (Figure 5A), when assuming a uniform placement of mutations along the length of a branch. We therefore introduce a new algorithm to account for time-heterogeneous mutation rates (details in Section), which can be shown to be much more accurate than when assuming uniform mutation placement (Figure 5A).

**Fig. 5.**
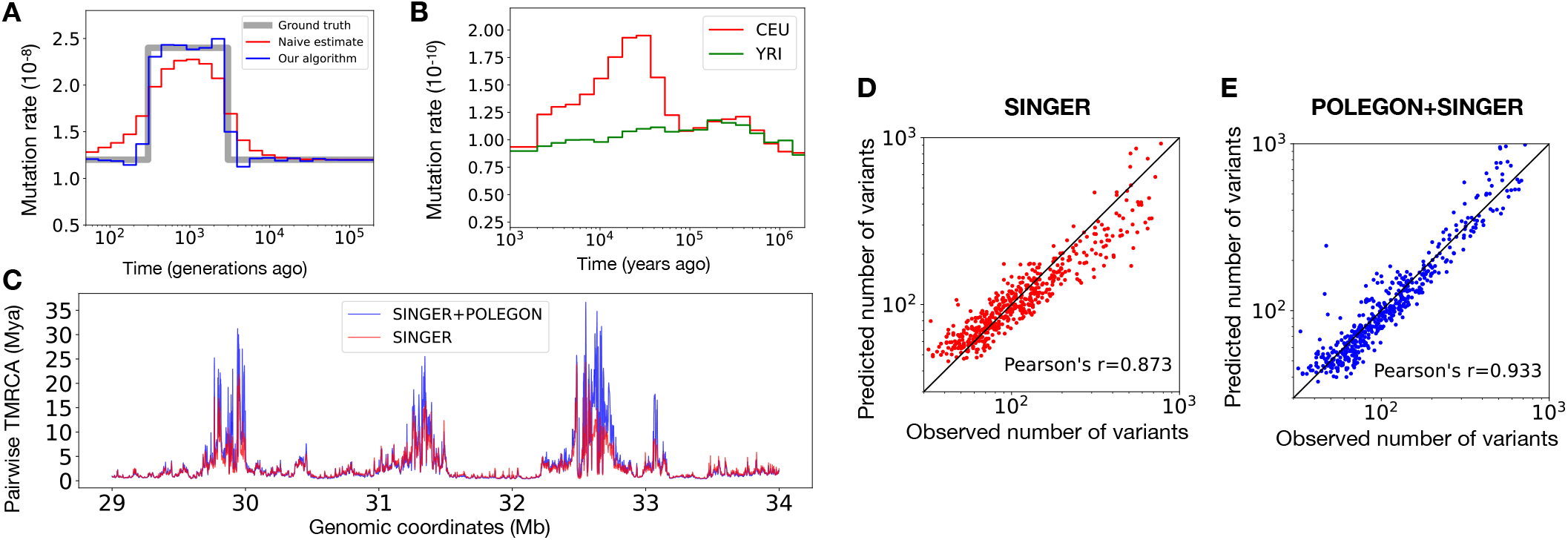
Mutation rate trajectory inference and ancient coalescence times at HLA inferred with SINGER+POLEGON. (A) Performance of mutation rate trajectory inference when assuming uniform placement (red) and using the proposed iterative algorithm (blue); (B) Inferred mutation rate trajectory for TCC → TTC in CEU and YRI, with our new algorithm; (C) The estimated average pairwise TMRCA in HLA region using SINGER (red) compared to SINGER+POLEGON (blue); (D) The number of observed variants versus predicted from inferred ARGs with SINGER in 10kb windows; (E) The number of observed variants versus predicted from inferred ARGs with POLEGON+SINGER in 10kb windows.

We have incorporated this new mutation rate trajectory estimation to the inferred genome-wide genealogies with SINGER+POLEGON in CEU and YRI and recover the signal of elevated TCC to TTC mutations specific to CEU (Figure 5B) and absent in YRI. As expected, using the uniform placement of mutations on branches leads to underestimation of mutation rate variation (Figure 5A). Our estimates of the timing of elevated mutation rates agrees well with those previously reported in (12). However, we note that (12) inferred a somewhat larger difference in the mutation rate of TCC to TTC mutations between modern Europeans and Africans. Previous literature (41) argued that the rate difference is not clearly observed in rare variants and suggested a potential recent loss of the mutation rate modifier. The difference between our estimates and those of (12) is likely due to the artifact induced by uniform placement of mutation on branches in Relate, similarly to that observed in Figure 5A. As noted earlier, we here assume a constant aggregate mutation rate (across all sub-types), which the ARG rescaling relies on, allowing us only to infer temporal changes in the relative mutation rates of different sub-types.

### Coalescence times in the HLA region

The HLA locus has been shown to harbor trans-species polymorphisms hypothesized to be under strong balancing selection (42). The presence of trans-species polymorphism would imply coalescence times in this region that are older than the divergence time between humans and chimpanzees, and (16) indeed shows this to be the case.

In (16) the mutation rate was assumed to be constant, which ignores the heterogeneity of mutation rate along the genome. Here we reanalyze the HLA data with POLEGON on the SINGER topologies (16) with a mutation map specific to the HLA region estimated with Roulette (43). The methods used to extract and process the mutation map can be found in *SI Appendix, SI Text*, Section D. The updated estimates are older than the the previous estimates from SINGER (Figure 5C), with some alleles having average pairwise coalescence time older than 30Mya, consistent with previous findings (44–48). However, we note that these ancient coalescence times estimates inevitably rely on short regions with elevated polymorphisms, which might provide noisy estimates. Also, the human mutation rate estimates used in POLEGON become less reliable when going beyond the divergence time of humans and other primates. As additional evidence, we also estimate phylogenetic gene trees on HLA allele sequences from multiple primate species, which identify trans-species polymorphisms between human and Old World Monkeys in the same regions as inferred by the ARG (*SI Appendix, SI Text*, Section D, *SI Appendix, SI Text*, Figure S6).

To validate that the estimates from POLEGON+SINGER indeed make more sense than using SINGER alone, we compared the observed mutation density versus the predicted mutation density from the inferred ARG (Figure 5D, E). With inferred genealogies, the predicted mutation density in a window is defined as the integral of the product of local tree branch length and local mutation rate over the window, divided by the window length. With 10kb genome windows, the mutation density predictions from using POLEGON+SINGER clearly correlate better than using SINGER alone, which suggests better inference quality (Figure 5D, E). Notably, there is a slight underestimation bias in SINGER with the higher mutation density regions, but it is largely corrected with SINGER+POLEGON. The bias in SINGER is expected as it uses a standard exponential coalescence prior that will tend to shrink the estimates towards zero.

## Discussion

In this article, we introduced POLEGON, a novel approach for estimating branch lengths in ancestral recombination graphs. POLEGON infers branch lengths and population size history given a fixed ARG topology using an uninformative uniform prior. The use of an uninformative prior provides increased accuracy and flexibility when the populations studied are not panmictic populations of constant size. We note that although the current version assumes the infinite-sites model, the model can potentially accommodate a finite-site model after redefining the likelihood of node ages and the rescaling step.

We applied POLEGON to SINGER-inferred genealogies for CEU and YRI populations from the 1000 Genomes Projects and estimated the demographic history of these groups to re-evaluate the evidence for a severe bottleneck in the YRI (33). Our results largely agreed with previous findings, and we did not find evidence of a strong bottleneck. Instead, consistent with other methods, we inferred a much milder bottleneck shared by CEU and YRI.

We introduced a new way of inferring temporal mutation rate patterns that it is not biased by the assumption of uniform mutation placement on branches. Analyses of the CEU and YRI data confirmed the observation of a TCC to TTC pulse in the CEU but not YRI. While the timing of the pulse agrees well with the estimates of (12), the difference in rates between CEU and YRI before and after the pulse are smaller than those reported by (12), which assume uniform placement of mutation on branches. Third, we re-dated the coalescence times in the ARG of the HLA region from (16) with POLEGON, while accounting for mutation rate heterogeneity using a mutation map inferred by Roulette (43). Our estimates suggest coalescence times as old as 30 million years in multiple different areas of the HLA region.

There are a few limitations to our work. Firstly, the demographic model we considered include only single population size changes in history and clean splits, and we have not investigated more complex models that include processes such as migration and admixture. Inferring node ages accurately in the presence of these factors may require an accurate reconstruction of ARG topology incorporating these processes, in addition to an accurate dating methods. However, we note that the assumption of exchangeability among lineages in the standard coalescent model, and assumed in all common methods for estimating ARGs, corresponds to an uninformative prior on the tree topology, i.e. a prior that is not informed by population designation of leaf nodes.

Secondly, the current methodology is based on the assumption of a constant aggregate mutation rate from all subtypes (e.g., defined by the triplet sequence context of the mutated site). Whether this is a reasonable assumption at all time scales remains to be investigated. If the pattern of temporal mutation rate variation has been inferred, then it is straightforward to incorporate into the MCMC algorithm and the subsequent ARG rescaling. It would perhaps make more sense to incorporate temporal mutation rate variation into the ARG estimation algorithm itself, rather than just at the branch length estimation step, but that is beyond the scope of this work focusing on branch length estimation for a given ARG topology. Of course, any dating methods will perform worse with lower mutation to recombination ratio, both due to the increasing difficulty of inferring ARG topology from mutation data and from the lack of branch length information from sparser mutation counts (Figure S2).

Thirdly, even more accurate estimates of coalescence times could potentially be obtained by using an estimated demographic history as prior for the node age estimation, similarly to the approach applied by (12). However, the use of an noninformative prior can provide unbiased (or at least less biased) estimates in regions with selection, as illustrated in our example for the HLA locus. Furthermore, it might also facilitate other downstream inferences by decoupling the estimates of coalescence times from the parametric assumptions regarding population history.

Finally, we note that the accuracy of coalescence time estimation is limited by the quality of the the topology inference. As such, future development of better methods for estimation topologies, can help improve coalescence time estimation.

## Materials and Methods

### The MCMC algorithm

To sample the node ages in the ARG (or equivalently, the converted DAG), we perform MCMC with a Metropolis-Hastings algorithm for each of the *N* nodes in the ARG.

#### Algorithm 1

MCMC for node ages in Ancestral Recombination Graph

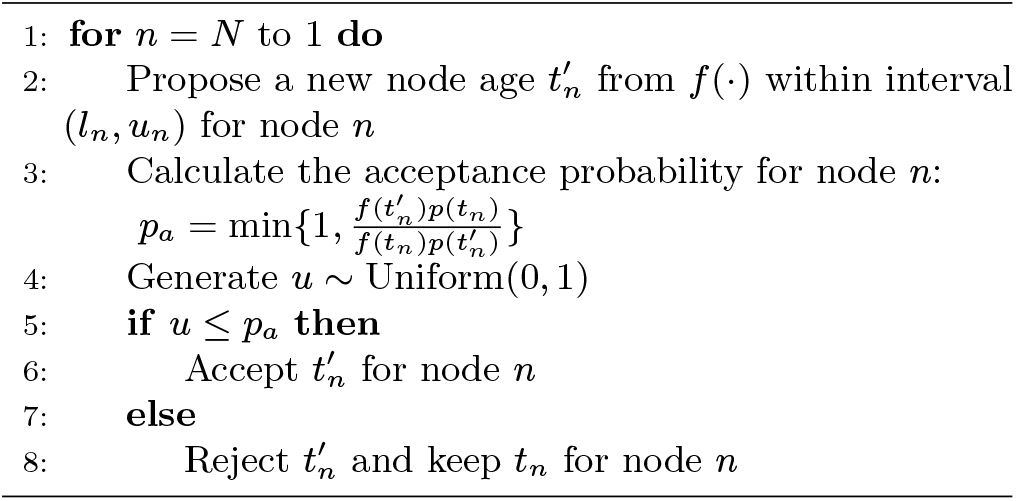

Here, *f* (·) is the proposal function which determines how we propose new ages in the interval (*l*_*n*_, *u*_*n*_). When the upper bound, *u*_*n*_, is not infinity, we simply propose uniformly at random in the interval (*l*_*n*_, *u*_*n*_).

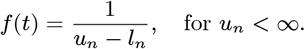

This results in the following acceptance probability:

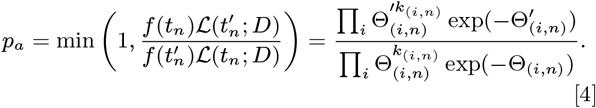

However, for the root node, *u*_*n*_ = ∞ and an alternative approach is needed. We then let *f* (·) equal an exponential distribution with a small rate parameter, *λ* (default at *λ* = 0.1 in POLEGON):

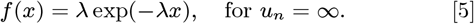

The exponential distribution approaches an improper uniform on [0, ∞) as *λ* → 0. Now the acceptance probability is:

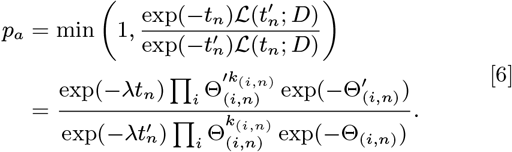

We also note that we chose the order of node age updates to be the descending order of the node ages, so that when a node age is proposed, then the ages of its parent nodes will have already been updated. This leads to more efficient mixing compared to randomly choosing new nodes to update. The full details of the MCMC can be found in *SI Appendix, SI Text*, Section A.

### Population size history estimation

To infer demography, we first need to define change points in time when the population size (or equivalently, coalescence rate) can change. In this paper we chose 30 windows log-uniformly distributed from 100 to 200,000 generations ago (generation time by default at 28 years for human). We simply extract the empirical pairwise coalescence distribution from the ARG, and obtain an empirical survival function *S*(*t*), i.e. the proportion of coalescence times larger than *t*.

Assume the time grid is (0, *t*_1_, *t*_2_, …, *t*_*K*_, ∞), and the coalescence rates are (*λ*_0_, *λ*_1_, …, *λ*_*K*_). The survival function should satisfy:

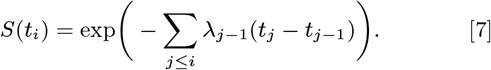

The coalescence rates can be solved by noticing:

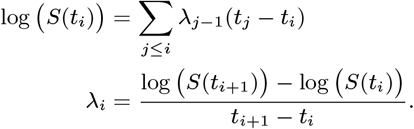

The coalescence rates are subsequently convert them into an estimate of the population size history.

### Estimation of temporal patterns of mutation rate variation

Here we introduce an algorithm for estimating temporal mutation rate variation from genealogies, inspired by (38). Assume the time grid is (0, *t*_1_, *t*_2_, …, *t*_*K*_, ∞) and the mutation rates are *µ* = (*µ*_0_, *µ*_1_, …, *µ*_*K*_) in the time intervals, we can define a design matrix **B**, whose entry *B*_*ij*_ stands for the overlap of ARG branch *i* with time interval (*t*_*j*_, *t*_*j*+1_):

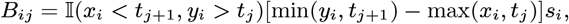

where (*x*_*i*_, *y*_*i*_) are the times of lower and upper node of branch *i*, and *s*_*i*_ is the spatial span of the branch *i*. Assume we observe **k** = (*k*_0_, *k*_1_, …, *k*_*m*_) mutations on all branches, we can formulate the mutation rate estimation as a optimization problem with the following log-likelihood (after dropping the constant terms):

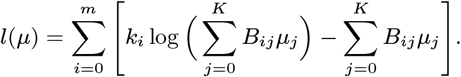

As the gradient can be analytically calculated, standard first order optimizer can be used to solve this with the analytical gradient.

## Data availability

POLEGON is available at https://github.com/YunDeng98/POLEGON.git. The ARG samples from SINGER+POLEGON are available at: https://zenodo.org/records/14675005, https://zenodo.org/records/14674978, https://zenodo.org/records/14676049, https://zenodo.org/records/14676128, https://zenodo.org/records/14676158.

## ACKNOWLEDGMENTS

We thank Drew DeHaas and Kaiyuan Li for testing the software. WSD thanks Peter Ralph and Kameron Decker Harris for feedback on an early version of the mutation rate inference procedure. This research is supported in part by NIH grants R35GM153400, R56-HG013117 and R01-HG013117. WSD was supported in part by a Fellowship in Understanding Dynamic and Multi-scale Systems from the James S. McDonnell Foundation.

## Notes

### Competing Interest Statement

The authors have declared no competing interest.

### Summary of Updates

"Mutation subtypes" clarified, the change is at line 785 in the Discussion section on the current limitation of the method.

