## Supplementary Information for "A General Framework for Branch Length Estimation in Ancestral Recombination Graphs" (from another version of this preprint: version 2, 2025-08-14)

**This PDF file includes:**

Supporting text
Figs. S1 to S7
Tables S1 to S2
SI References

### Supporting Information Text

#### 1. Method details

**A. The full algorithm.** In the first step, we first use the variant data and the input mutation rate to determine the $N_e$ so that $4N_e\mu$ matches the nucleotide diversity $\pi$. We then re-scale node ages by $2N_e$ generations such that underflow or overflow problems are unlikely to occur. If a spatial mutation map is used, rather than a fixed mutation rate, we use the mean mutation rate to carry out the same operation.

We use the the Metropolis-Hastings MCMC algorithm as described in Section 1. In our default settings, the first 100 iterations are used as burn-in and we then sample with thinning in intervals of 10 iterations. In each iteration, new ages are proposed iteratively for all the nodes in the ARG.

POLEGON can provide both the posterior average and posterior samples for each node age. For the posterior average, we first calculate the average node age among the posterior samples. These average ages will conform to the DAG constraints too. We then carry out the ARG rescaling operation on the ARG with the new ages assigned, for 5 iterations by default. For the posterior samples, we similarly perform ARG rescaling on each sample separately by sampling no node ages (also for 5 iterations by default).

The users can also set their own parameters following the documentation at https://github.com/YunDeng98/POLEGON.git, including thinning interval, number of samples, ARG rescaling iterations, etc.

**B. The coalescent prior induced by conditional uniformity.** Our MCMC algorithm makes proposals based on the likelihood function using a uniform prior, conditional on all other node ages that may constrain the age of the node currently being updated. This induces an implicit prior of coalescence times when integrating over all possible ages of the other nodes in the graph. This implicitly induced prior corresponds to the joint distribution of inter-coalescence waiting times being uniform. To verify this, we ran the MCMC algorithm with no mutation data on single coalescent trees each with $c+1$ leaf nodes (hence with $c$ coalescence events), and generated samples from our MCMC algorithm. The sample distribution of inter-coalescence waiting times should reflect the induced prior on coalescence times. (We note that we have to set an upper bound as we cannot simulate from a uniform prior.) More precisely, the ordered inter-coalescence waiting time proportions (i.e., waiting times divided by the TMRCA) in a random tree sampled from the MCMC should follow a uniform Dirichlet distribution $\mathrm{Dir}(1,\ldots,1)$ on a $(c-1)$-dimensional simplex. This is because if there are $c$ i.i.d exponential random variables $X_1\ldots,X_c$ with the same rate parameter $\lambda$, then the joint distribution for the proportions $(\frac{X_1}{S},\ldots,\frac{X_c}{S})$, where $S=\sum_{i=1}^{c}X_i$, is the uniform distribution on the $(c-1)$-dimensional simplex, and the improper uniform distribution can be seen as the limit of $\lambda\to 0$. This implies that the $k$-th inter-coalescence waiting time divided by the TMRCA should follow the $\mathrm{Beta}(1,c-1)$ distribution, for all $1\le k\le c$. As illustrated in Figure S1A-D for the case of 5 leaf nodes without recombination, the MCMC output fits the $\mathrm{Beta}(1,3)$ distribution very well for all inter-coalescence waiting time proportions, showing that the algorithm indeed induces a uniform joint distribution for inter-coalescence waiting times.

We note that, this also holds more generally for a full ARG with $C$ coalescence events (or equivalently, $C$ coalescence nodes). In the full ARG with recombination, the joint distribution of all inter-coalescence waiting time proportions should also follow a uniform Dirichlet distribution, and the $k$-th inter-coalescence waiting time proportion should follow a $\mathrm{Beta}(1,C-1)$ distribution. To verify this, we carry out simple coalescent simulations again with 5 leaf nodes but with recombination. We then pick ARGs with different number of coalescences, and plot the aggregate distribution of all the inter-coalescence waiting time proportions in the ARG. Again, the results agree well with the predicted $\mathrm{Beta}(1,C-1)$ distribution (Figure S1E-H).

We note that there are connections between these two cases. We used simulations to show that when the inter-coalescence waiting time proportions follow the uniform Dirichlet distribution in the simulated ARGs, then in the marginal trees, the inter-coalescence waiting time proportions also follow the uniform Dirichlet distribution (Figure S1I-M).

**C. Posterior distribution calibration with ARG rescaling.** In this section we will show why using improper uniform prior and ARG rescaling can help achieve the correct posterior distribution even when the true prior is unknown, using both theory and simulation benchmarks.

**Pairwise case:** We first consider a simple model of pairwise coalescent, where the mutation rate is $\theta/2$ on either branch, and the population size is constant at $1/\lambda$. Under Kingman's coalescent the distribution for the coalescent time $t$ is exponential distribution:

$$p(t)=\lambda\exp(-\lambda t).$$

The number of observed mutations $k$ is a Poisson distribution conditional on the coalescent time $t$:

$$p(k|t)=\frac{(\theta t)^k}{k!}\exp(-\theta t).$$

The unconditional distribution for the number of mutations in the pairwise coalescent tree follows a geometric distribution, where the success probability is the probability that a mutation event happens before the coalescent event:

$$p(k)=\frac{\lambda}{\lambda+\theta}\left(\frac{\theta}{\lambda+\theta}\right)^k.$$

If we know the true prior distribution of $t$, then the posterior distribution of $t$ is a Gamma distribution:

$$p(t|k) \propto p(t)p(k|t) = \Gamma(k+1, \theta+\lambda).$$

However, if we use an improper uniform prior, then we hfave a mis-specified posterior distribution:

$$\tilde{p}(t|k) \propto p(k|t) = \Gamma(k+1, \theta).$$

Now suppose we perform ARG rescaling in the pairwise coalescent case. Then, the mutation counts between $[s, s+\Delta s]$, $g(s)\Delta s$, is:

$$\begin{aligned}
g(s)\Delta s &= \int_s^\infty \sum_{k=1}^\infty \frac{k\Delta s}{t}\tilde{p}(t|k)p(k)dt \\
&= \Delta s \int_s^\infty \sum_{k=1}^\infty \frac{k}{t}\left[\frac{\theta^{k+1}t^k}{k!}\exp(-\theta t)\right]\frac{\lambda}{\lambda+\theta}(\frac{\theta}{\lambda+\theta})^k dt \\
&= \Delta s \int_s^\infty \left[\sum_{k=1}^\infty \frac{(\frac{\theta^2 t}{\lambda+\theta})^{k-1}}{(k-1)!}\right]\exp(-\theta t)\frac{\lambda\theta^3}{(\theta+\lambda)^2}dt \\
&= \Delta s \int_s^\infty \exp\left(\frac{\theta^2 t}{\lambda+\theta}\right)\exp(-\theta t)\frac{\lambda\theta^3}{(\theta+\lambda)^2}dt \\
&= \Delta s \frac{\theta^2}{\lambda+\theta}\exp\left(\frac{-\lambda\theta s}{\lambda+\theta}\right).
\end{aligned}$$

In the first equality, $\frac{k\Delta s}{t}$ denotes the contribution of mutation counts to the window $[s, s+\Delta s]$ assuming a uniform mutation mapping when the pairwise coalescence time is $t$, the integration of $t$ starts from $s$ because the contribution is non-zero only when the pairwise coalescence time is larger than $s$, and we integrate over both the pairwise coalescence time $t$ and the observed number of mutations $k$.

The prior corresponding to the mis-specified posterior $\tilde{p}(t|k)$ of pairwise coalescent times is:

$$\begin{aligned}
\tilde{p}(t) &= \sum_{k=0}^\infty \tilde{p}(t|k)p(k) \\
&= \sum_{k=0}^\infty \frac{\theta^{k+1}t^k}{k!}\exp(-\theta t)\frac{\lambda}{\lambda+\theta}\left(\frac{\theta}{\lambda+\theta}\right)^k \\
&= \frac{\lambda\theta}{\lambda+\theta}\exp(-\theta t)\sum_{k=0}^\infty \frac{(\frac{\theta^2 t}{\lambda+\theta})^k}{k!} \\
&= \frac{\lambda\theta}{\lambda+\theta}\exp(-\theta t)\exp\left(\frac{\theta^2 t}{\lambda+\theta}\right) \\
&= \frac{\lambda\theta}{\lambda+\theta}\exp\left(\frac{-\lambda\theta t}{\lambda+\theta}\right).
\end{aligned}$$

This indicates an inflation factor of $\frac{\lambda+\theta}{\theta}$ for the pairwise coalescence time, as the true distribution should be of rate $\lambda$. The inflation is less severe with higher mutation rates, which makes sense since the mis-specified prior should matter less with more data.

Now we investigate the effects of ARG rescaling. We first compute the expected branch length between $[s, s+\Delta s]$, $f(s)\Delta s$, as:

$$\begin{aligned}
f(s)\Delta s &= \Delta s \int_s^\infty \tilde{p}(t)dt \\
&= \Delta s \exp\left(\frac{-\lambda\theta s}{\lambda+\theta}\right).
\end{aligned}$$

Again, the integration starts from $s$ because if the coalescence time is lower than $s$, there will be no contribution of branch length to the window $[s, s+\Delta s]$. This means that the “effective mutation rate" in the posterior samples (assuming the improper uniform prior) at time $s$ is:

$$m(s) = \frac{g(s)}{f(s)} = \theta\frac{\theta}{\lambda+\theta}.$$

Because the true mutation rate is $\theta$, we need to apply a rescaling factor of $\frac{\theta}{\lambda+\theta}$ to calibrate the mutation clock. Rescaling the uncalibrated posterior distribution $\tilde{p}(t|k)$ with a factor of $\frac{\theta}{\lambda+\theta}$ then induces the correct posterior $p(t|k)$. It also translates the uncalibrated pairwise coalescent time distribution $\tilde{p}(t)$ to the correct distribution $p(t)$. So even without knowledge of $\lambda$, use of an improper uniform prior with posterior rescaling induces the correct posterior distribution that would have been obtained using the correct prior, regardless of the value of $\lambda$.

**General ARGs:** Similar theory is less tractable for general ARGs, so we choose instead to use simulations to show that POLEGON samples with recalibration provide a good characterization of the posterior distribution that would have been obtained, if the natural prior induced by the generative model had been used (Figure 2).

##### D. Varying ratio of mutation to recombination rates.

We next evaluated the performance of POLEGON with different ratios of mutation rate ($\mu$) to recombination rate ($r$). We simulated with both $\mu/r = 4$ and $\mu/r = 0.25$ for the same three demography models used in the main text (Section A), and run POLEGON with true ARG topology. Again, we observe that the rank plots are close to a uniform distribution after ARG rescaling and the inferred pairwise TMRCA distributions matches the ground truth well (Figure S2).

We note that the posterior distributions tend to be more over-dispersed with lower mutation rates than with higher mutation rates. This is likely due to the ARG rescaling step, where more mutations lead to be more accurate rescaling, and fewer mutations can lead to noisy rescaling, which explains the bigger variance in the posterior distribution.

##### E. Mutation rate heterogeneity.

If there is spatial variation in mutation rates and a mutation rate map is available, then $\Theta_{(i,n)}$ will have to be calculated by integrating over the mutation rate map, instead of using the formula in Eq. (1). Suppose the mutation rate is $\mu(x)$ at position $x$ in the genome, then the value of $\Theta_{(i,n)}$ for a branch $(i, n)$, which spans spatially from $x_l$ to $x_r$ in the genome should be defined as:

$$\Theta_{(i,n)} = (t_i - t_n) \int_{x_l}^{x_r} \mu(x)dx.$$

If the mutation rate map is fine-scale (such as base-pair resolution) then this integration can be computationally expensive to calculate. However, we note that we only have to store the cumulative sum of mutation rates, $S(x)$, at the positions of recombination breakpoints along the genome, because the integrals will always be defined between these breakpoints. The integral can then be evaluated quickly:

$$\int_{x_l}^{x_r} \mu(x)dx = S(x_r) - S(x_l).$$

In this work, we use the mutation map inferred by Roulette (1). However, there are two issues with this mutation map: first, the values in the inferred map are only proportional to the per-bp per-generation mutation rate, so we re-scale the rates globally to ensure that the genome-wide average rate is $1.2 \times 10^{-8}$; second, the map does not cover every region in the genome, so for the gaps, we treat them as having the genome-wide average mutation rate of $1.2 \times 10^{-8}$.

#### 2. Simulation benchmarks

##### A. Simulation settings.

The CEU and the YRI model used in the simulations were taken from (2), which used SMC++ (3) on the 1000 Genomes Project Phase 3 data (4). The coalescent simulation was performed with msprime (5), with $\mu = r = 1.2 \times 10^{-8}$ on 10Mb chunks for 10 replicates.

##### B. Demography inference with more pairs.

Compared to methods like PSMC or MSMC, which can only practically use one or few pairs of sequences, ARG-based methods like Relate and POLEGON can utilize more information as it effectively involves many more pairs of sequences. These pairs are not independent but still provide substantially more information, and estimates based on more pairs should lead to more stable results. To demonstrate this, we extracted one, eight, and a hundred pairs from the terminal sample of ARG from SINGER (re-dated with POLEGON) in 10 simulation replicates under the CEU model. Then we perform demography inference and compare these estimates with the ground truth (Figure S3). It is clear that with the usage of more pairs, the estimates get more consistent across replicates and closer to the ground truth.

##### C. Clean split model.

In this work we mainly tested the performance of POLEGON on branch length estimation and demography inference under panmictic models, but we also simulated a simple structured model: a clean split of two populations (both with $N_e = 10,000$) which split 10,000 generations ago. See Figure S4.

##### D. Simulations with temporally varying mutation rates.

Coalescent simulation with temporally varying mutations rate is not available in msprime (5), so we designed the following simulation pipeline. Firstly, we simulated the whole-genome genealogies with $N_e = 10,000$ and $r = 1 \times 10^{-8}$. We then computed the Poisson rate on each persisting branch. Suppose the mutation rate trajectory over time is $\mu(t)$, and assume a branch $(c, p)$ with child node at time $t_c$ and parent node $t_p$, which spatially spans from $x_l$ to $x_r$. Then, the rate parameter can be calculated as:

$$\Theta_{(c,p)} = (x_r - x_l) \int_{t_c}^{t_p} \mu(t)dt.$$

Here the rate function $\mu(t)$ is chosen to be $2.5 \times 10^{-8}$ from 300 to $3,000$ generations ago and $1 \times 10^{-8}$ for the rest (black in Figure 5A).

We then simulate the mutation counts on each branch from a Poisson distribution with the corresponding Poisson rate. The simulated genealogies and mutation counts for each persisting branch are the input data for Figure 5A and the mutation rate trajectory inference algorithm.

**E. Settings when running Relate, SINGER, and POLEGON.** For simulation benchmarks, we compared the performance of Relate, SINGER and SINGER+POLEGON, for all the demographic models mentioned above. We ran SINGER and Relate with $\mu = r = 1.2 \times 10^{-8}$. For the MCMC setting in SINGER, we used "-n=100, -thin=20". We additionally re-estimate branch length in Relate with "EstimatePopulationSize.sh". We then ran POLEGON with default settings on every MCMC sample.

**F. Dating tsinfer topologies.** To assess the performance of POLEGON on dating ARG topologies other than ground truth and SINGER inferred topologies, we choose to use tsinfer to infer topology and then use tsdate and POLEGON to estimate branch lengths. We use the same benchmark as in the main text to assess the quality of dating from these two methods.

POLEGON produces more accurate dating results under the CEU and YRI models, and similar results under the constant size model (Figure S5). This is likely because tsdate uses a coalescent prior derived from the standard coalescent (constant size), and thus is able to work well under the constant size model. The performance of tsdate is worse than POLEGON when the underlying model is not a constant size model, due to prior mis-specification.

Tsinfer tends to infer topologies with polytomies whereas SINGER never does so, and we note that special methods for dating should probably be applied to such polytomies (which POLEGON probably does not cover). We also note that the error, in general, are much larger for Tsinfer topologies than for SINGER-estimated ARG topologies (Figure 3). This suggests that the performance of the dating methods are fundamentally limited by the inference quality of the topology.

**G. Quantification of demography inference accuracy.** To quantitatively evaluate the accuracy of the population size history estimates from different methods, we chose to use the L2 distance between the ground truth coalescence rate and the estimated coalescence rates in varying time windows (2kya-2Mya). The reason why the L2 distance is calculated in the space of coalescence rate, rather than population size, is to avoid the error being dominated by the large population size estimates obtained for the recent past, which can be common for methods like MSMC2. The results are summarized in Table S1.

#### 3. Real data analysis

**A. Application of SINGER+POLEGON to data from the 1000 Genomes Project.** We downloaded the 1000 Genomes Project high coverage (30X) VCF files from https://www.internationalgenome.org/data-portal/data-collection/30x-grch38, and extracted variant calls for the YRI and CEU for autosomes (chr1-22). For each of these VCF files, we removed all the non-SNP variants (structural variants, multi-allelic sites, etc) and polarized SNPs with respect to the primate ancestral genome downloaded from https://ftp.ensembl.org/pub/release-109/fasta/ancestral_alleles/. For the SINGER analysis, the mutation rate was set to $1.2 \times 10^{-8}$ globally, and the population size was set to $2 \times 10^4$ and $1.5 \times 10^4$ for YRI and CEU, respectively, following the guidelines in the SINGER documentation (https://github.com/popgenmethods/SINGER) to match their respective nucleotide diversities. Since the VCF files are polarized with respect to an ancestral genome, we used the polarized version of SINGER with the "-polar 0.99" flag.

Similarly to (6), we used $15,000$ MCMC iterations with the first $5,000$ iterations as burn-in and 100 iterations as the thinning interval length, which led to a total of 100 posterior ARG samples for each population. After obtaining ARG samples from SINGER, we ran POLEGON with the default parameters and other procedures as described in Section A.

**B. Application of Relate and MSMC2 to the 1000 Genomes Project data.** We ran Relate on the same dataset as in the section above (Section A), with $\mu = r = 1.2 \times 10^{-8}$. We subsequently used "EstimatePopulationSize.sh" to re-estimate branch lengths and population size histories. After obtaining the coalescence rate estimates for each chromosome, we calculated the weighted average of coalescence rates (based on chromosome length) to obtain the final demography model.

We similarly applied MSMC2 to all within-individual haplotype pairs and used the default setting for time discretization, optimization iterations, etc. The genome-wide coalescence rates are obtained as weighted averages according to chromosome length, similarly to the procedures for Relate.

**C. Mutation signatures across populations.** We used the primate ancestral sequence to determine the triplet mutation context before and after the mutation. Variants whose ancestral allele states were uncertain (marked as "·" or "N" in the inferred ancestral genome fasta file) were left out from the mutation signature analysis. We then extracted the TTC to TCC mutations and then used the mutation rate trajectory estimation algorithm mentioned in Section 1 of the main text.

**D. Re-dating HLA with POLEGON.** We used the mutation map inferred by Roulette (1), and used POLEGON to re-date the ARGs in the HLA region inferred in (6). We note that the mutation rates inferred by Roulette do not cover every base pair in the genome, so we chose to fill the gaps with the genome-wide average rate ($1.2 \times 10^{-8}$). We ran POLEGON on the inferred ARG samples from (6) with the default settings.

To further validate our results on trans-species polymorphism, we employed phylogenetic methods to construct gene trees using HLA sequences from humans and other primates. We obtained HLA locus sequences from the IPD-IMGT/HLA database (https://www.ebi.ac.uk/ipd/mhc/), which includes data from both human and non-human primates. Sequences were grouped by HLA gene, and multiple sequence alignments were generated using MAFFT. Gene trees were then inferred with IQ-TREE, using automatic model selection to ensure appropriate substitution models.

In accordance with previous results (7), we observed clear evidence of trans-species polymorphism in several HLA genes, where some human alleles are more closely related to alleles from other primates than to other human alleles. For instance, in *HLA-DQA1* and *HLA-DQB1*, we identified trans-species polymorphism between humans and, minimally, Old World monkeys (Figure S6), which indicates coalescence time above 30 Mya (the assumed split time to Old World Monkeys (8)). To validate our gene tree inferences and guard against false positives, we also analyzed genes known to have low levels of trans-species polymorphism as negative control. As expected, gene trees for *HLA-E* and *HLA-F* closely matched the species tree, with human sequences forming a distinct monophyletic clade (Figure S6).

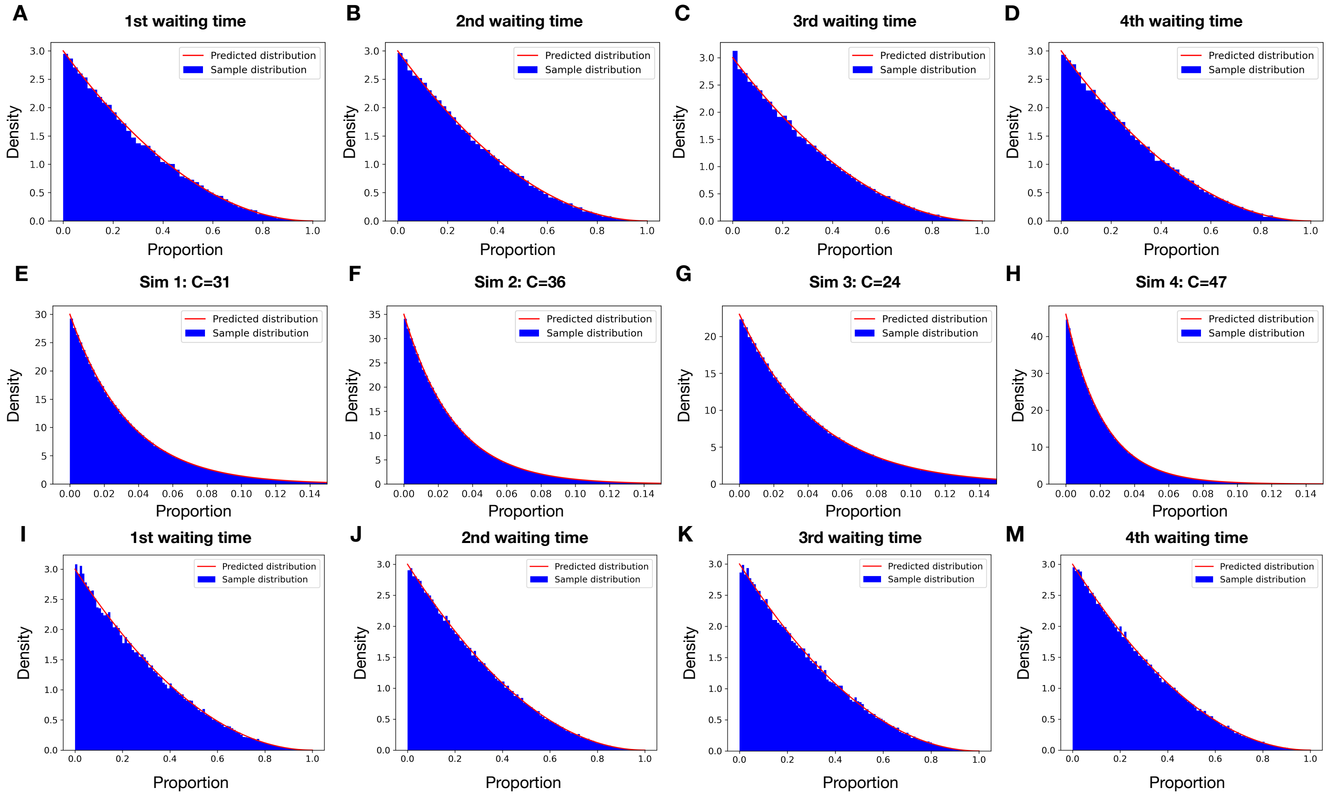


**Fig. S1.** The proportion of inter-coalescence waiting time follows the predicted Beta distributions, for both a single coalescent tree (A-D) and a full ARG (E-H). When examining marginal trees from an ARG whose inter-coalescence waiting time proportions follow the Beta$(1, C-1)$ distribution, all inter-coalescence waiting time proportions in the trees follow the expected Beta$(1, c-1)$ distribution (I-M), where $c$ and $C$ correspond to the number of coalescence events in a marginal coalescent tree and in the full ARG, respectively.

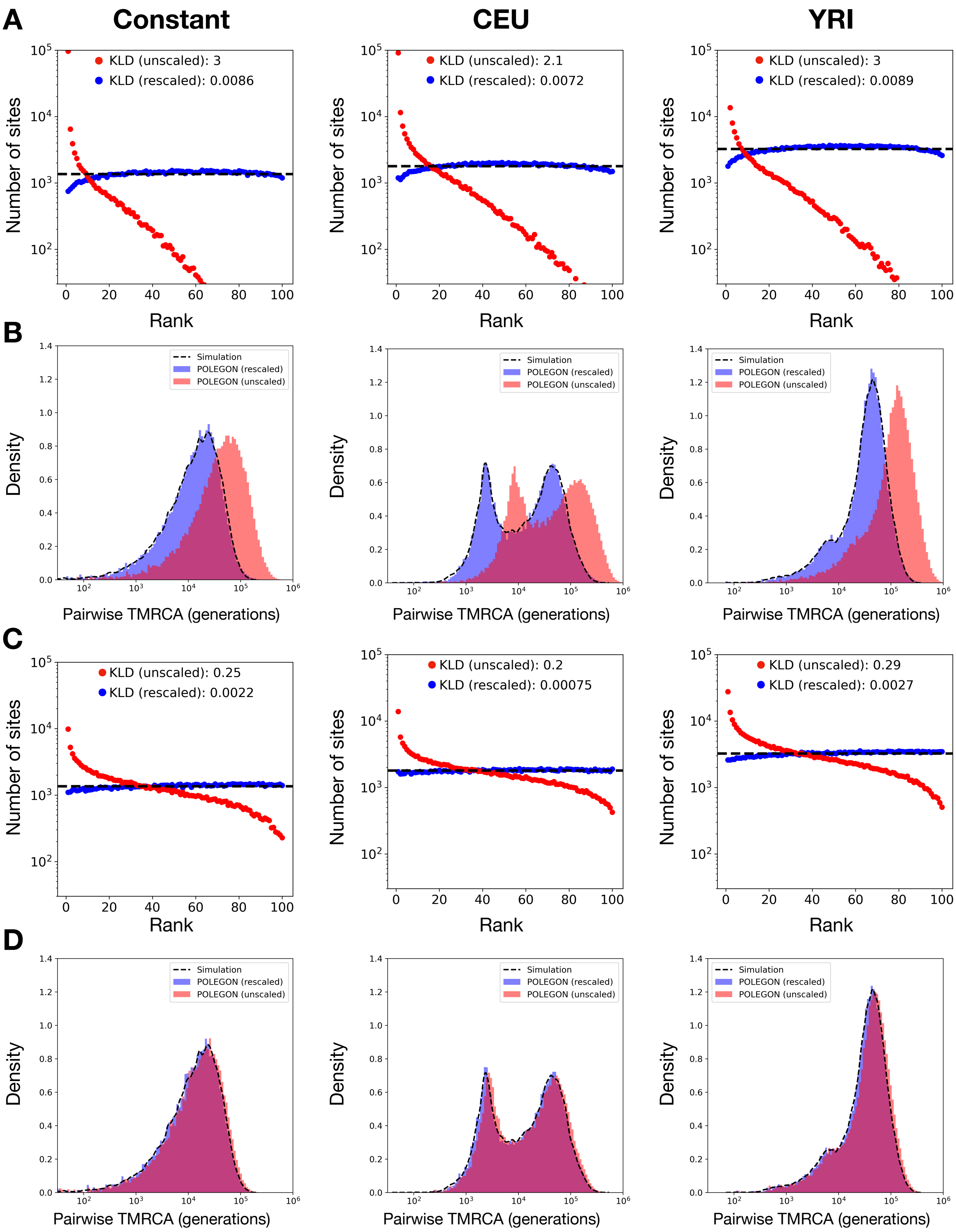


**Fig. S2.** The MCMC sampling performance with $\mu/r = 0.25$ (A, B) and $\mu/r = 4$ (C, D). The performance was benchmarked through rank plots (A, C) and the comparison of pairwise TMRCA distribution to ground truth (B, D).

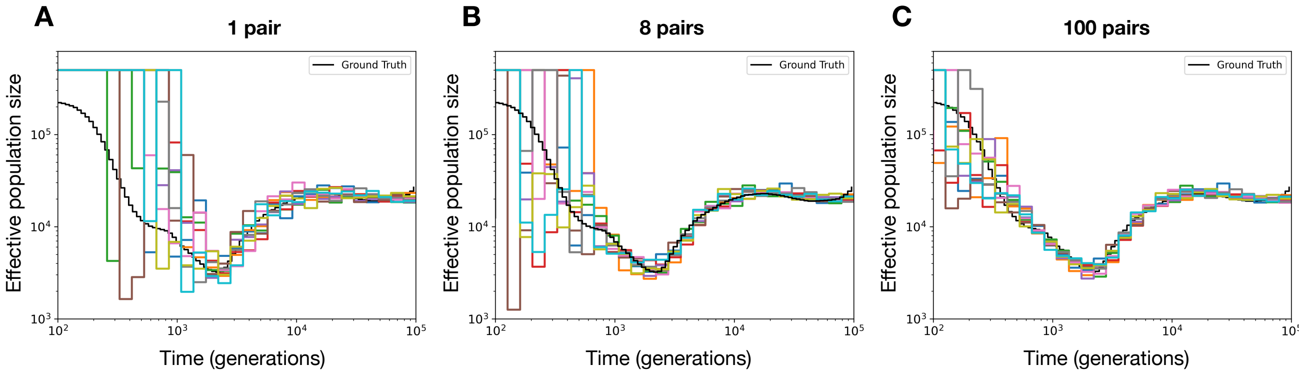


**Fig. S3.** Demography inference results (colored lines) from 10 simulation replicates compared with the ground truth (black line), using one (A), eight (B), and a hundred pairs (C) extracted from the inferred ARG. Each colored line is the inferred demography from one terminal sample of ARG from SINGER (re-dated with POLEGON).

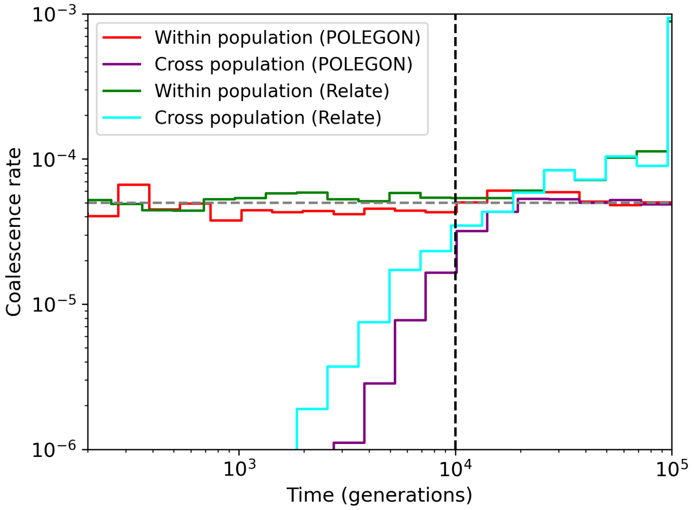


**Fig. S4.** The coalescence rate inference under the clean split model with POLEGON+SINGER and Relate. The split time is indicated by the vertical black dashed line and the true within coalescence rate is indicated by the horizontal gray dashed line. Coalescence rates are computed both within populations and across populations. Both methods infer that the cross-population coalescence rate decrease rapidly compared to the within-population coalescence rate after the divergence time.

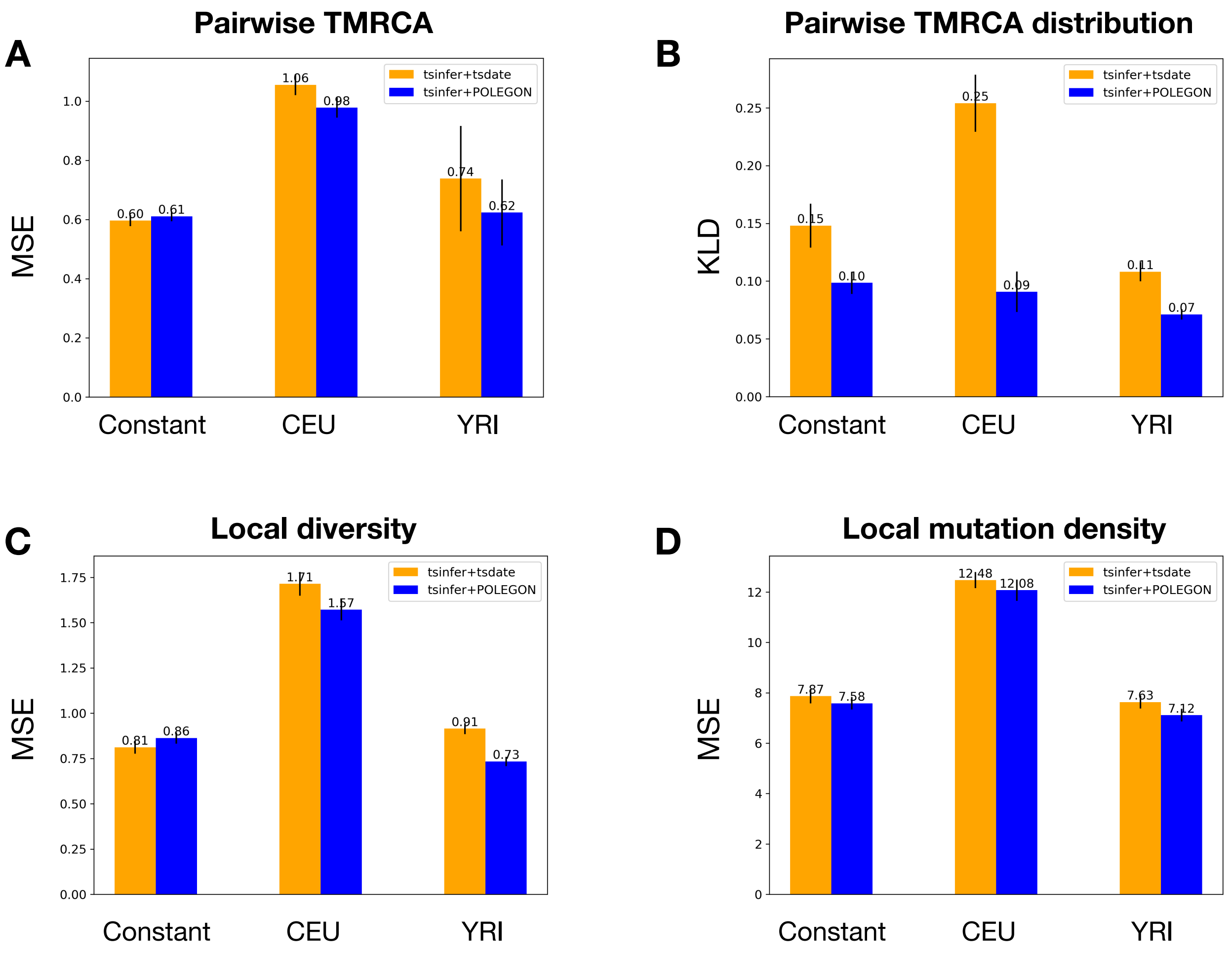


**Fig. S5.** The inference accuracy comparison between tsdate (orange) and POLEGON (blue) on ARG topology inferred by tsinfer.

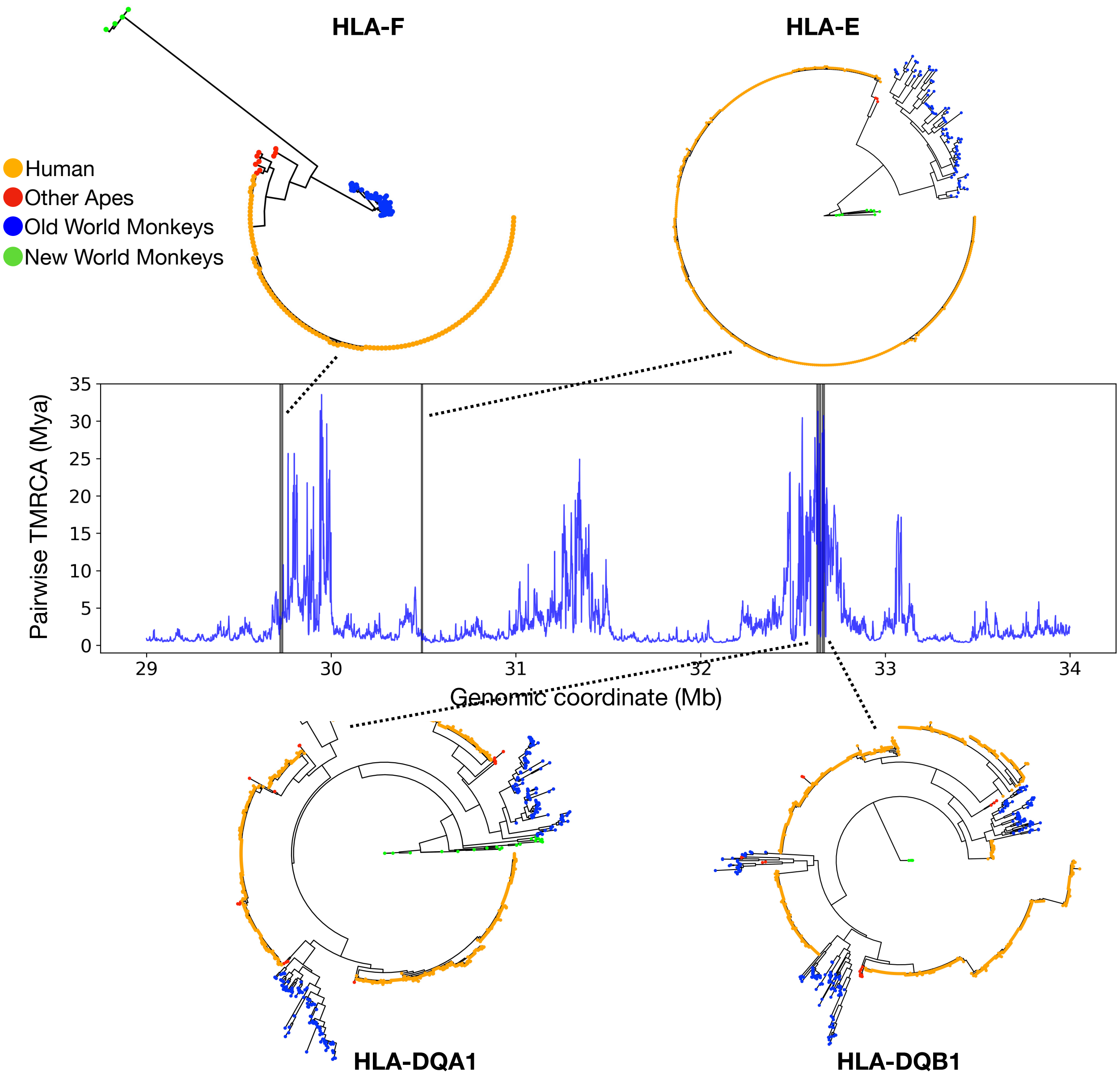


**Fig. S6.** Gene trees of primate species inferred with IQ-TREE at different HLA loci.

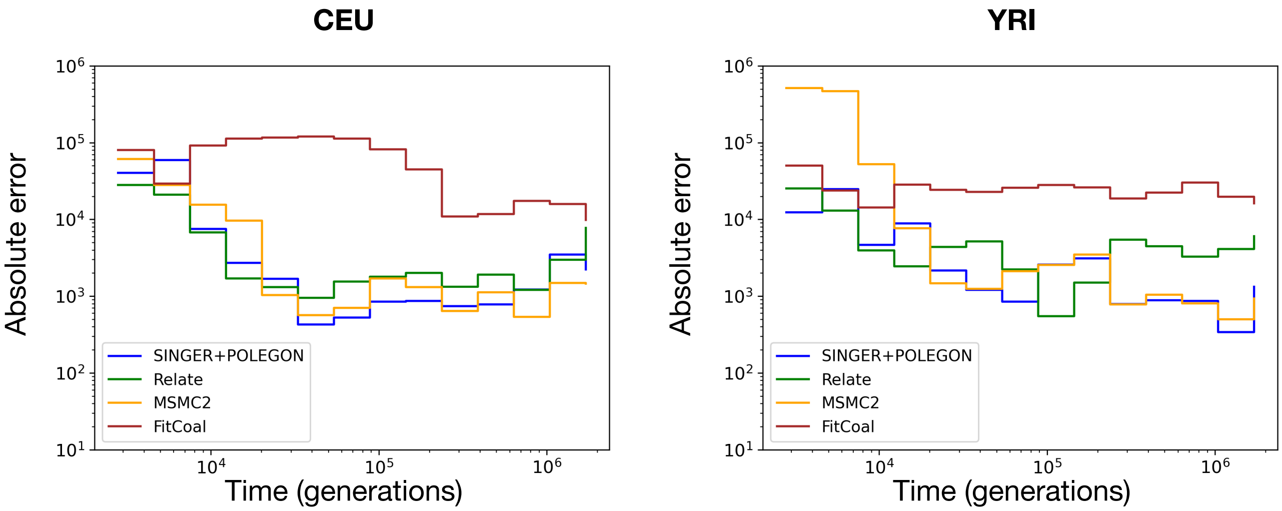


**Fig. S7.** The absolute error in effective population size estimation at different times under the CEU and YRI model, with different demography inference methods.

| Method | CEU | YRI |
|---|---|---|
| SINGER+POLEGON | $2.49 \times 10^{-3}$ | $6.05 \times 10^{-4}$ |
| Relate | $4.65 \times 10^{-3}$ | $3.22 \times 10^{-3}$ |
| FitCoal | $5.68 \times 10^{-2}$ | $7.11 \times 10^{-3}$ |
| MSMC2 | $2.70 \times 10^{-3}$ | $6.82 \times 10^{-4}$ |

**Table S1. Comparison of demographic inference methods under CEU and YRI models.**

| | Without rescaling | With recaling |
|---|---|---|
| Constant | 0.0460 | 0.0007 |
| CEU | 0.2946 | 0.0016 |
| YRI | 0.1186 | 0.0004 |

**Table S2. KL divergence between the pairwise TMRCA distribution from simulated ARGs and POLEGON inference on simulated ARGs, with and without the rescaling step.**
